# Multi-omics analyses reveal the effect of DNA methylation on iridoid glycosides biosynthesis in *Rehmannia glutinosa*

**DOI:** 10.1101/2025.06.11.659110

**Authors:** Tianyu Dong, Yajie Du, Tingting Huang, Jiuchang Su, Qingxiang Yang, Jie Guo, Peilei Chen, Jingjing Xing, Hongying Duan

## Abstract

*Rehmannia glutinosa* roots produce a group of lipophilic bioactive components known as iridoid glycosides. However, the molecular mechanisms by which DNA methylation regulates the biosynthesis of iridoid glycosides in *R. glutinosa* remain unknown. Herein, the development of *R. glutinosa* roots and the content of iridoid glycosides in the Wenxian region were significantly higher than those in Xinxiang. Low methylation level contributed to the accumulation of iridoid glycosides and the expression of related enzyme genes. Demethylation promoted both roots growth and development, as well as the accumulation of iridoid glycosides. Up-regulated *RgALDH13*, *RgHDR1*, *RgG10H4*, *RgDXR1*, *RgG10H3*, and *RgUPD1*, along with transcription factors (TFs), form the regulatory network for the biosynthesis of iridoid glycosides. Furthermore, the primary active region of the *RgG10H4* promoter is located in the -164 bp region, where the RgMYB2 protein specifically binds to the TAACCA motif in the *RgG10H4* promoter. Collectively, low levels of DNA methylation enhance the expression of core genes, followed by inducing the accumulation of iridoid glycosides, which suggests that *RgMYB2*-*RgG10H4* plays a positive role in this process. These findings will contribute to a deeper understanding of the role of DNA methylation in the accumulation of iridoid glycosides.

**Highlight:** Low levels of DNA methylation contribute to the accumulation of iridoid glycosides and the expression of key enzyme genes in *R. glutinosa*.

## Introduction

Plant secondary metabolites (PSMs) are small molecular compounds that are not essential for primary metabolic processes, but hold substantial economic, industrial, and medicinal significance (Hartmann, 2007; Saslis-Lagoudakis et al., 2012). The PSMs are primarily composed of alkaloids, flavonoids, terpenes, tannins, phenolic acids, quinones, steroids, and phenylpropanes. Previous studies have shown that the accumulation of PSMs is influenced by environmental factors such as temperature, humidity, light exposure, and CO2 concentrations (Yang et al., 2018; Li et al., 2020; Wang et al., 2020; Jamloki et al., 2021; Jan et al., 2021).

Environmental changes frequently induce modifications in the DNA methylation status of plant genomes (Thiebaut et al., 2019; Chang et al., 2020). The processes of DNA methylation and demethylation collaboratively regulate the expression of essential enzyme genes and transcription factors (TFs), ultimately impacting the biosynthetic pathways of PSMs (Yang et al., 2021; Ferrandino et al., 2023; Zang et al., 2023). In *Eleutherococcus senticosus*, moderate water deficiency decreased global DNA methylation levels, including those in the promoter regions of *EsFPS*, *EsSS*, and *EsSE*. This demethylation enabled the re-binding of EsMYB-r1 to the *EsFPS* promoter region, thereby enhancing gene expression and leading to the accumulation of saponins (Wang et al., 2024). The genomic DNA of American Ginseng underwent complete demethylation at low temperatures, resulting in an increased expression level of the ginsenoside biosynthesis gene *PqDDS* and elevated ginsenoside accumulation (Hao et al., 2020). In tea plants, the methylation levels of CG and CHG sites across the genome increased, while those of CHH sites decreased during the summer. Higher temperatures triggered the expression levels of genes for DNA methyltransferases and demethylases, which subsequently upregulated the expression of enzymes and TFs involved in the metabolic pathways of flavonoids and theanine (Han et al., 2024).

Up to now, 5-azacytidine (5-azaC), a DNA methylation inhibitor, has been utilized to explore the correlation between DNA methylation and the accumulation of PSMs. Treatment with varying concentrations of 5-azaC suppressed the growth and development of *Pogostemon cablin*, while significantly increasing the levels of patchouli alcohol and pogostone (Luo et al., 2021). In *Salvia miltiorrhiza*, the tanshinone content in hairy roots increased by 1.5 to 5 times following 5-azaC treatment, likely due to the demethylation of 51 cytosines in the promoter region of the gene encoding copolymer diphosphate synthetase (Yang et al., 2022). The genomic DNA of *Bixa orellana* was demethylated following treatment 5-azaC. The expression levels of *ABATH* and *CCD4* were down-regulated, while *ALDH3H1* was up-regulated, collectively promoting accumulation of bixin in leaves (Faria et al., 2020). In strawberries, the 5-azaC treatment resulted in increased levels of proanthocyanins, (epi)afzelechin, and epi(catechin), as well as elevated expression of key enzyme genes, thereby promoting fruit ripening (Martínez-Rivas et al., 2022). Similarly, the demethylation of the citrus genome promoted chlorophyll degradation, enhanced carotenoid biosynthesis, and increased limonene content. This process influenced peel color and fruit ripening by modulating the expression levels of *CLH*, *PAO*, *RCCR*, *ZEP*, and *NCED* (Chen et al., 2024). Transcriptomic analysis revealed that differentially expressed genes (DEGs) were significantly enriched in pathways associated with photosynthesis, flavonoid synthesis, and glutathione metabolism in grape leaves subjected to 5-azaC treatment. Key enzyme genes, along with MYB, C2H2, and bHLH TFs, constitute an interaction network that regulates the biosynthesis of PSMs (Jia et al., 2020). In a word, although the relationship between DNA methylation and the accumulation of PSMs has been investigated, the underlying molecular mechanisms through which DNA methylation regulates PSMs have yet to be elucidated.

*Rehmannia glutinosa* is a dual-purpose crop valued for its industrial and dietary applications, particularly due to its high concentration of iridoid glycosides, which hold significant economic importance (Dong et al., 2022; Zhou et al., 2023; Li et al., 2024). In this study, we found that the DNA methylation levels of *Rehmannia glutinosa* in the Wenxian region are relatively low, while the content of iridoid glycosides and the expression levels of key enzyme genes are markedly high. DNA demethylation led to an increased accumulation of iridoid glycosides, enhanced expression of key enzyme genes, and a rise in the abundance of the crucial protein RgG10H4. Furthermore, the functional region of the *RgG10H4* promoter was cloned and characterized, revealing that the RgMYB2 specifically binds to the promoter of *RgG10H4*. Overall, our findings elucidate the role of DNA methylation in the accumulation of iridoid glycosides in *Rehmannia glutinosa*, offering novel insights into the epigenetic development and utilization of crops with applications in industry, food, and medicine.

## Materials and methods

### Cultivation and treatment of R. glutinosa

The roots of *R. glutinosa* “Jinjiu” were separately planted at Henan Normal University and the Wenxian Agricultural Science Research Institute in early April. The *R. glutinosa* roots were harvested at distinct developmental stages, designated as stage E (characterized by fleshy, cylindrical roots observed in early June), stage I (marked by root expansion occurring in the middle of September), and stage M (distinguished by spindle-shaped roots evident in early December). Additionally, a subset of 30 *R. glutinosa* root tubers at stage I underwent treatment with 50 μМ 5-azaC, with three independent replicates conducted. The aforementioned harvested roots were stored at - 80°C for preservation.

### Quantification of iridoid glycosides content

As previous study described (Dong et al., 2022), the roots of *R. glutinosa* were dried to a constant weight at 50 °C in an oven, followed by pulverization of the dried sample. Subsequently, 1 g of the powdered sample was combined with 10 mL 70% ethanol solution, transferred into a 100 mL conical flask, and subjected to ultrasonic filtration. The resulting filtrate was then filtered and agitated in a 50 mL volumetric flask, followed by dilution of 2.5 mL of the filtrate to a final volume of 50 mL.

For the assessment of iridoid glycosides content, catalpol was employed as the reference standard. Specifically, 1 mL of the test solution was mixed with 2 mL of 1 M HCl in 10 test tubes, and the mixture was subjected to heating in a water bath at 90°C for 15 minutes, followed by cooling to room temperature for an additional 15 minutes. Subsequently, 0.5 mL of dinitrophenylhydrazine ethanol solution and 30 mL of 70% ethanol solution were added to the reaction mixture. The resulting solution was allowed to stand at room temperature for 1 h, and its absorbance was measured at 463 nm using a spectrophotometer.

### Widely targeted metabolomics analysis

Based on the methods of previous studies (Chen et al., 2013; Wang et al., 2019), the *R. glutinosa* roots were freeze-dried using a specialized machine and then ground into a fine powder with a grinder. A quantity of 50 mg of the powder was dissolved in 1.2 mL of a 70% methanol extract, followed by rotational mixing for 30 s at 30 min intervals, for a total of 5 repetitions. After centrifugation at 12,000 rpm for 3 min, the supernatant was carefully pipetted and filtered through a 0.22 µm Hydrophilic Teflon syringe filter (SCAA-104, ANPEL, Shanghai, China) before being stored in an injection bottle. Subsequently, ultra-high-performance liquid chromatography (UPLC) coupled with tandem mass spectrometry (MS/MS) analysis was performed using a SHIMADZU Nexera X2 UPLC system and an Applied Biosystems 4500 QTRAP MS/MS instrument to analyze the sample.

Metabolites were characterized by comparing their m/z values, retention times, and fragmentation patterns with standards from a self-compiled database (MetWare) (Schymanski et al., 2014). The precursor ions of interest were selectively isolated using a quadrupole, which eliminated interference from ions of different molecular weights. The relative abundance of each compound was determined by evaluating the signal intensity of the characteristic fragment Q3. After acquiring metabolite profiles from various samples, the mass spectrometry peak areas of all compounds were integrated. The multi-quant software was employed to process the mass spectrometry peaks and normalize identical metabolic peaks across different samples. The relative abundance of metabolites was quantified based on the area under each peak (Fraga et al., 2010). In addition, differentially abundant metabolites (DAMs) were identified based on fold change values of ≥ 1.2 or ≤ 0.83, with statistical significance set at *P* < 0.05. The DAMs, along with their associated metabolic pathways, were annotated using the Kyoto Encyclopedia of Genes and Genomes (KEGG) database.

### RNA-sequencing and qRT-PCR

Total DNA was extracted using the RNAprep Pure Plant kit, and RNA integrity was assessed with the Agilent Bioanalyzer 2100 (Agilent Technologies, CA, USA). Subsequently, mRNA enrichment was performed using Oligo(dT) beads to target polyadenylated mRNA molecules. The synthesis of first-strand cDNA and second-strand cDNA was accomplished following a standardized procedure. After quality assessment of the library, sequencing was conducted using the Illumina NovaSeq 6000 system to generate 150 bp paired-end reads. The raw sequencing data were then filtered using fastp software to obtain clean reads (Chen et al., 2018). These clean reads were assembled using Trinity software (Grabherr et al., 2011). Transcript or gene expression levels were quantified using FPKM (Fragments Per Kilobase of transcript per million fragments mapped) (Wei et al., 2024). Differential expression analysis was performed using the DESeq2 R package, with P-values adjusted using the Benjamini–Hochberg method to control the false discovery rate. Differentially expressed genes (DEGs) were identified based on the criteria of |log2(Fold Change)| ≥ 1 and adjusted *P*-value ≤ 0.05. Quantitative real-time PCR (qRT-PCR) was performed using the LightCycler 96 fluorescence quantitative PCR system, with *RgTIP41* designated as the internal reference gene. The primer sequences are listed in Supplementary Table 1. The reaction was prepared in accordance with the instructions provided by the SYBR Green MasterMix kit (Vazyme, Nanjing, China). Target gene expression levels were determined utilizing the 2^^-ΔΔCt^ method (Wang et al., 2021).

### Methylation-sensitive amplification polymorphism

The methylation-sensitive amplified polymorphism (MSAP) protocol, as outlined by Dong et al. (2022), was utilized in this study. Genomic DNA underwent EcoRI/MspӀ (M) and EcoRI/HpaII (H) digestion, followed by ligation with EcoRI and MspI-HpaII adapters at a temperature of 16°C for 15 h. The resulting products were then utilized for MSAP amplification. The MSAP pre-amplification product served as the template for MSAP selective amplification. Subsequently, the DNA methylation banding patterns were categorized into three groups based on the presence or absence of bands in H and M: 1) no methylation (class I), assigned to the presence of bands in both H and M; 2) DNA hemi-methylation (class II), assigned to the presence of bands solely in H; and 3) DNA full methylation (class III), assigned to the presence of bands exclusively in M. Additionally, the MSAP percentage was calculated using the formula (II + III) / (I + II + III) × 100, and the percentage of fully methylated levels was determined as III / (I + II + III) × 100.

### iTRAQ quantitative proteomics

As described in a previous study (Chen et al., 2021), the protein extract was subjected to 12% SDS-PAGE electrophoresis, followed by reduction, cysteine blocking, and digestion using the FASP technique (Jacek et al., 2009). Subsequently, in accordance with the guidelines of the iTRAQ 8-plex kit, the protein samples were iTRAQ-labeled, resuspended, washed for 8 minutes, and then desiccated using a vacuum freeze dryer. The desiccated peptide samples were then resuspended and analyzed via RPLC-MS/MS. The primary protein data were scrutinized using the Paragon algorithm to correlate with digital transcriptome expression profiles. The experimental tandem mass spectrometry data were matched with theoretical data to obtain protein identification results. Using *Arabidopsis* as the reference organism, reliable proteins were identified based on sequence similarity and transcriptome sequences. Differentially expressed proteins (fold change > 1.2 or < 0.83, *P* < 0.05) under 5-azaC treatment were subsequently isolated from the pool of reliable proteins.

### Subcellular localization of RgG10H4

The coding sequence (CDS) of *RgG10H4*, which lacks a stop codon, was integrated into the Super1300-GFP vector, resulting in the creation of the Super1300-RgG10H4-GFP recombinant vector. The empty vector and the Super1300-*RgG10H4*-GFP vector were then introduced into *Agrobacterium tumefaciens* GV3101 and combined with mCherry to infect 4-week-old tobacco leaves. Subsequently, images were captured using a confocal laser scanning microscope (Leica, Germany).

### Cloning and activity analysis of RgG10H4 promoter

The *RgG10H4* promoter sequence was cloned using the fusion primer and nested integrated PCR (FPNI-PCR) method (Wang et al., 2011). Genomic DNA obtained from *R. glutinosa* root was subjected to extraction, followed by specific amplification utilizing fusion primers in the initial PCR round. The products were then recovered and purified, followed by a second round of PCR employing nested primers for further specific amplification. Detection was achieved through agarose gel electrophoresis, and subsequent verification was conducted through DNA sequencing.

Based on the functional regions of *cis*-acting elements, the 5’ truncated promoter fragments were designed. Subsequently, the pCAMBIA 1301 vector and the *RgG10H4* promoter truncation fragment were individually cleaved using *HindIII* and *BglII* restriction endonucleases. The 5’ truncated *RgG10H4* promoter fragment replaced the *CaMV35S* promoter through homologous recombination, resulting in the generation of GUS fusion plant expression vectors G10H4-P1 (1014 bp), G10H4-P2 (690 bp), G10H4-P3 (356 bp), and G10H4-P4 (164 bp). As precious study described (Wang et al., 2012), the recombinant plasmids were introduced into the *Agrobacterium tumefaciens* strain GV3101 and subsequently infiltrated into 4-week-old tobacco leaves. The leaves underwent dark incubation overnight followed by placement in a greenhouse (16 h light/8 h dark cycle, 22°C) for 2 days. Finally, β-glucosidase activity in the tobacco leaves was measured using the β-glucosidase (β-GC) assay kit (Solarbio, BC2560, Beijing, China).

### Molecular docking and yeast one-hybrid (Y1H) assay

The interaction between the proteins RgMYB2, RgC2H2-2, and RgbZIP3 with the *RgG10H4* promoter was simulated using HDOCK v1.1 and PyMOL 2.5.0 software. The nucleotide sequence of the *RgG10H4* promoter, along with the amino acid sequences of RgMYB2, RgC2H2-2, and RgbZIP3, was separately input as receptors and ligands in the HDOCK module (http://hdock.phys.hust.edu.cn/) (Yan et al., 2020). Subsequently, molecular docking simulations were conducted using PyMOL 2.5.0 to select polar docking atoms and visualize the interactions between the proteins and nucleotides. The confidence score (CS) was utilized to assess the likelihood of binding between the two molecules, calculated as CS = 1.0 / [1.0+e^0.02*(docking^ ^score^ ^+150)^]. A CS value greater than 0.7 indicates a high probability of binding between the two molecules. The Y1H assay was carried out following the methodology (Dong et al., 2022).

Specifically, the full-length sequences of RgMYB2, RgC2H2-2, and RgbZIP3 were inserted into the pGADT7 vector at the EcoRI and BamHI restriction sites. The resulting recombinant plasmid was co-transformed into yeast Y1HGold along with pAbAi-*RgG10H4*. The pGADT-53 vector was used as a positive control. Subsequently, all transformed candidates were cultured on SD/-Ura/-Leu medium in the presence of either 0 or 200 ng/mL of Aureobasidin A (AbA) for a duration of 3 days.

### Statistical analysis

The statistical analysis of the data was conducted using SPSS 10.0 software, and graphical representations were generated with GraphPad Prism 5.0. Each experiment included three biological replicates, and the results are expressed as means ± standard deviation (SD) from three separate trials. The experimental data were evaluated using Student’s *t*-test, with significance indicated by different letters on the bars (*P* < 0.05).

## Results

### Morphology and iridoid glycosides content in roots of Rehmannia glutinosa

*Rehmannia glutinosa* produced in Wenxian region is renowned for its high quality. Here, the morphological characteristics and iridoid glycosides content in *Rehmannia glutinosa* root were determined. At E stage, the length and maximum cross-sectional diameter of *R. glutinosa* roots in Wenxian were significantly greater than those in Xinxiang. At I stage, there was no significant difference in root length between Wenxian and Xinxiang, while the maximum cross-sectional diameter was significantly larger in Wenxian than in Xinxiang. The results at M stage were contrary to those observed at the I stage (Figure 1A-C). Furthermore, the content of iridoid glycosides in *R. glutinosa* roots gradually increased from E to M stage (Figure 1D). Additionally, the iridoid glycosides content in the roots of Wenxian was significantly higher than that in Xinxiang at both I and M stages (Figure 1D).

**Figure 1.**
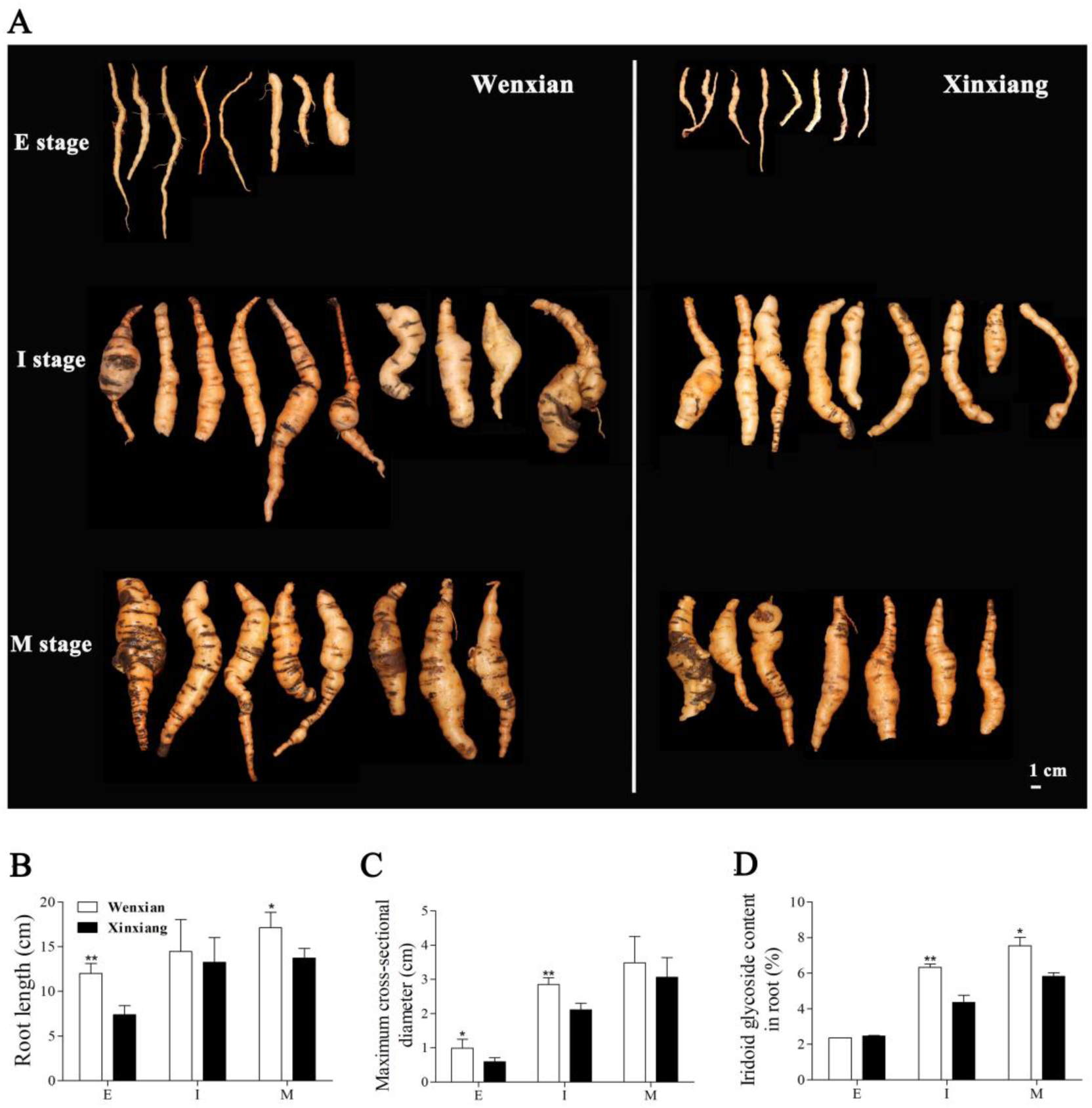
The morphology and iridoid glycosides content of *R. glutinosa* roots at E, I, and M stages in the producing areas of Wenxian and Xinxiang. (A) Morphological characteristics of *R. glutinosa* roots. (B), (C), and (D) The root length, maximum cross-sectional diameter, and iridoid glycosides content of *R. glutinosa* roots. The data are shown as the mean ± SD from three independent biological replicates (*, *P* < 0.05; **, *P* < 0.01; ***, *P* < 0.001; Student’s *t*-test).

### Analysis of differentially accumulated metabolites (DAMs)

Plant metabolomics was employed to explore the specialized metabolites in the roots of *R. glutinosa* at I stage. A total of 918 metabolites were detected and quantified, including 258 terpenoids, 242 phenolic acids, 145 alkaloids, 69 lignans and coumarins, 60 flavonoids, 13 quinones, 10 steroids, 4 tannins, and 117 others compounds (unknown classifications) (Figure S1; Table S2). The terpenoids comprised 103 monoterpenoids, 61 sesquiterpenoids, 60 triterpenes, 17 diterpenoids, 15 terpenes, and 2 triterpene saponins (Table S2). Principal component analysis (PCA) revealed that PC1 and PC2 accounted for 43.56% and 19.58% of the total variance, respectively (Figure S2). Further hierarchical clustering analysis (HCA) confirmed that the concentrations of most detected metabolites varied significantly between Wenxian and Xinxiang, based on the relative abundance of all annotated metabolites, which was consistent with the PCA results (Figure S3). Additionally, the correlation heatmap indicated a strong correlation among replicates within groups (Figure S4). Next, a total of 396 DAMs were identified, with 191 being up-regulated and 205 down-regulated in the treatment group (Wenxian) compared to the control group (Xinxiang) (Figure S5; Table S3). As expected, the HCA revealed significant differences in the accumulation of DAMs between Wenxian and Xinxiang (Figure S6). Among the DAMs, 122 terpenoids, 114 phenolic acids, 60 alkaloids, 29 lignans and coumarins, 10 flavonoids, 4 quinones, and 57 other compounds were identified (Figure 2A). The top ten up- and down-regulated DAMs between Wenxian and Xinxiang were primarily terpenoids, phenolic acids, and alkaloids, as determined by the analysis of fold change (FC) values (Figure 2B). Furthermore, the differentially accumulated monoterpenoids, terpenoids, diterpenoids, sesquiterpenoids, terpenes, and triterpene saponins were 45, 40, 4, 24, 7, and 2, respectively (Figure 2C). The identified DAMs were significantly enriched in monoterpenoid biosynthesis, phenylalanine metabolism, and tryptophan metabolism (Figure 2D). The differential abundance score (DA Score) for monoterpenoid biosynthesis exhibited an upward trend, indicating that the monoterpenoid metabolites in the roots of *R. glutinosa* from Wenxian were higher than those from Xinxiang (Figure 2E). In addition, 13 iridoid glycosides were identified among the monoterpenoids, of which 9 were up-regulated and 4 were down-regulated. The levels of catalpol, geniposide acid, geniposidic acid, adoxosidic acid, geniposide, gentiolactone, and ajugol, which are the primary iridoid glycosides in *R. glutinosa*, were higher in Wenxian than in Xinxiang (Table S4).

**Figure 2.**
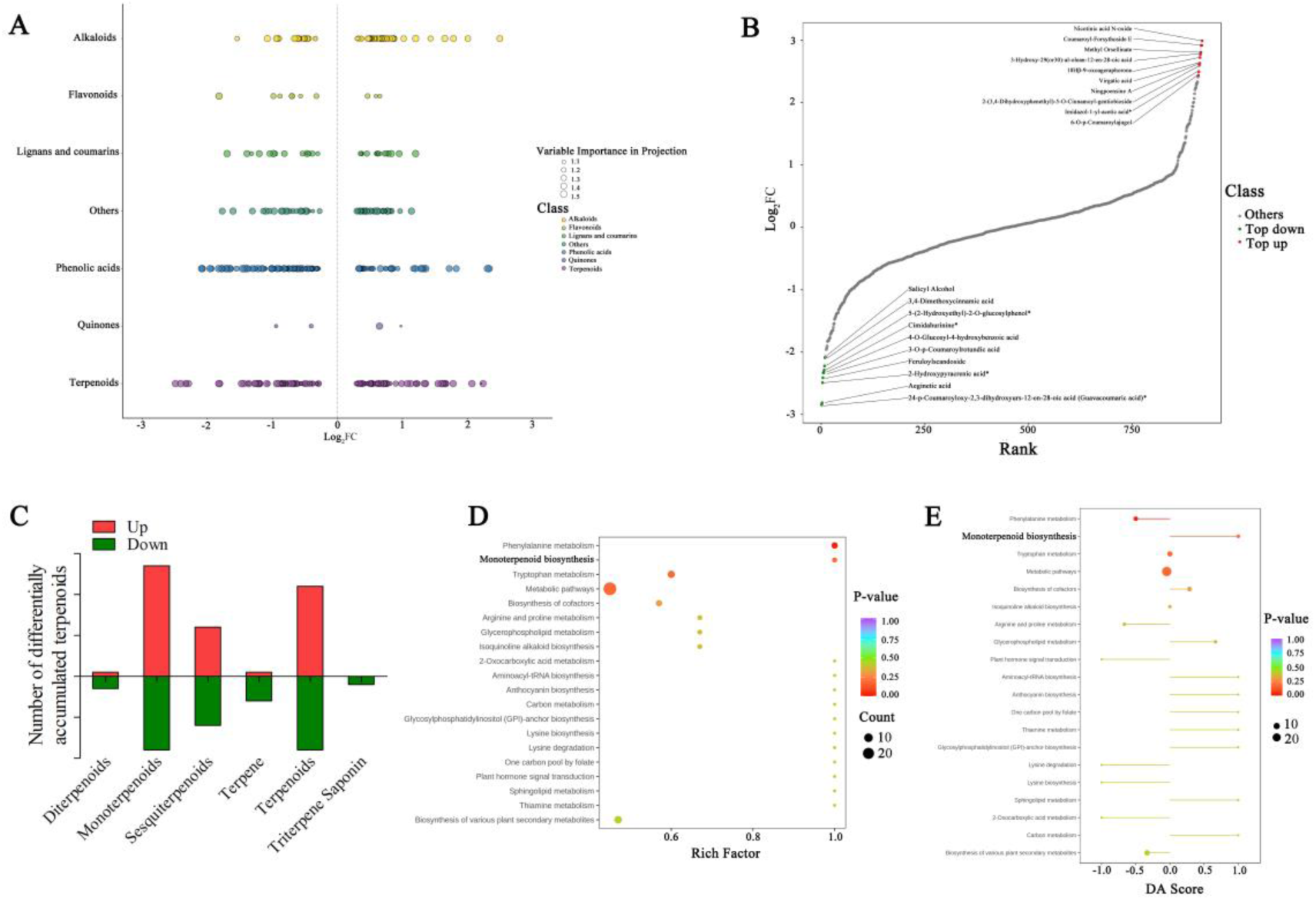
Analysis of differentially accumulated metabolites (DAMs) in *R. glutinosa* roots. (A) Scatter plot of DAMs. Color modules indicate various metabolites. (B) The top ten up- and down-regulated DAMs. (C) The differentially accumulated of terpenoids. (D) KEGG enrichment analysis of DAMs. The lower p-value indicates more significantly enriched pathways, and the point size indicates the number of DAMs. (E) Differential abundance of metabolic pathways. The DA Score reflects the overall changes in DAMs of KEGG pathways. 1.0 and -1.0 indicate up- and down-regulated of DAMs. The line length represents the absolute value of the DA Score.

### Analysis of differentially expressed genes (DEGs) involved in iridoid glycosides biosynthesis

The analysis of DAMs revealed that the levels of iridoid glycosides in Wenxian were significantly higher than those in Xinxiang. To gain molecular insights into the DAMs in the roots of *R. glutinosa*, transcriptome sequencing was performed on samples from Wenxian (treatment group) and Xinxiang (control group) at I stage. A total of 9,437 DEGs were identified, with 4,579 genes being up-regulated and 4,858 down-regulated (Figure S7). Clustering heatmap analysis indicated significant variations in the DEGs between Wenxian and Xinxiang (Figure S8). GO annotation of the DEGs indicated that the metabolic process was enriched in the biological process (BP) category, while catalytic activity was enriched in the molecular function (MF) category, both of which are closely related to terpenoid metabolism (Figure 3A). The KEGG analysis found that a substantial number of DEGs were enriched in monoterpenoid biosynthesis. Specifically, the expression levels of *Rg10HGO*, *RgMNR*, and *RgHDG* were significantly up- or down-regulated (Figure 3B). Further analysis of DEGs involved in iridoid glycoside biosynthesis identified 43 up-regulated DEGs and 10 down-regulated DEGs (Table S5). All terpene synthases (TPS), which are crucial enzymes in the production of various terpene structures, were significantly up-regulated. UDP-dependent glycosyltransferases (UGTs) catalyze the glycosylation process, which represents the final step in the biosynthesis of iridoid glycosides. The expression levels of most UGTs were significantly up-regulated (Figure 3C). Additionally, protein-protein interaction (PPI) predictions revealed that the genes responsible for iridoid glycoside biosynthesis interact with various enzymes, including O-methyltransferase, sorbitol dehydrogenase, catalase isozyme, and zeaxanthin epoxidase (Figure 3D; Table S6).

**Figure 3.**
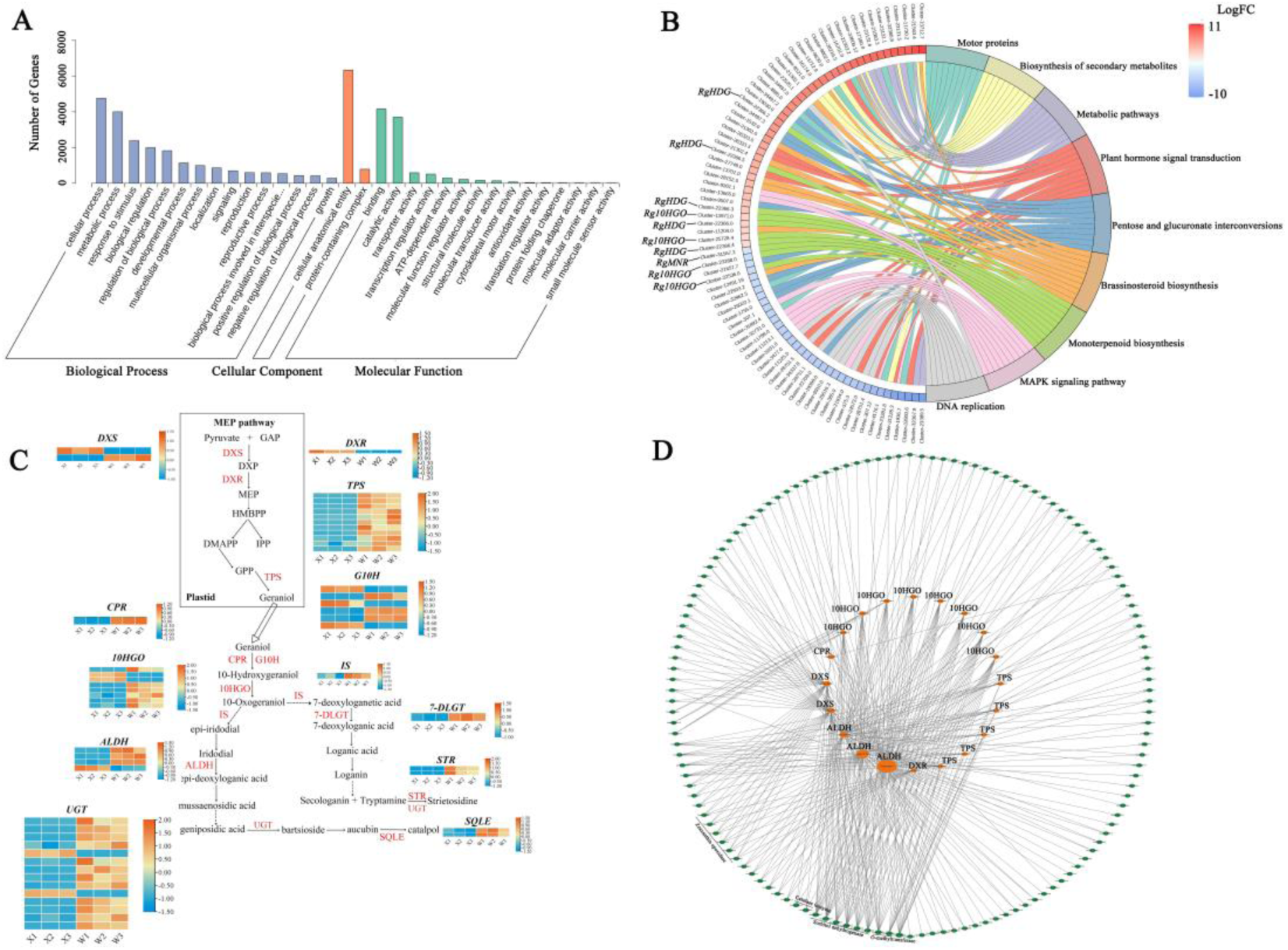
Analysis of differentially expressed genes (DEGs) in *R. glutinosa* roots. (A) The DEGs under the histogram of GO classification. (B) The chordplot of KEGG enrichment. On the right are the nine most significantly enriched KEGG pathways, and on the left are the 10 genes with the highest |logFC| in each pathway. Red and blue represent up- and down-regulated genes, and the color depth indicates the magnitude of |logFC|. (C) DEGs involved in the biosynthesis of iridoid glycosides. Red and blue represent up- and down-regulated genes. (D) Correlation network of genes involved in iridoid glycosides biosynthesis. The genes involved in the iridoid glycoside pathway are depicted in orange, while the genes that interact with them are depicted in green.

### Determination of genomic DNA methylation in R. glutinosa roots

Various growth environments lead to alterations in genomic DNA methylation, which in turn affect the accumulation of secondary metabolites. We hypothesized that the accumulation of iridoid glycosides and the expression of related genes in the roots of *R. glutinosa* from were significantly higher than those from Xinxiang, potentially due to varying levels of DNA methylation. Here, MSAP amplification was employed to detect the genome methylation ratio in roots of *R. glutinosa*. At the I stage of Xinxiang roots, 116 bands were analyzed, consisting of 55 unmethylated bands, 34 hemi-methylated bands, and 27 fully methylated bands. In contrast, a total of 80 bands were detected in Wenxian roots, which included 54 unmethylated bands, 15 hemi-methylated bands, and 11 fully methylated bands (Table 1, Figure S9). The fully methylated ratio of *R. glutinosa* root was 23.28 in Xinxiang and 13.75 in Wenxian County, while the MSAP ratio was 52.59 in Xinxiang and 32.50 in Wenxian (Table 1). The genomic methylation level of *R. glutinosa* root in Xinxiang was markedly greater than that in Wenxian. To further investigate this disparity, methyltransferase and demethyltransferase genes were identified from the transcriptomes of both Xinxiang and Wenxian. The findings revealed a notably higher expression level of the methyltransferase gene in Xinxiang compared to Wenxian (Figure S10A), whereas the expression level of the demethyltransferase gene was significantly lower in Xinxiang than in Wenxian (Figure S10B).

**Table 1.** The MSAP ratio of *R. glutinosa* root in Xinxiang and Wenxian at I stage.

| MSAP amplified type | I stage |  |
| --- | --- | --- |
|  | Xinxiang | Wenxian |
| Class I | 55 | 54 |
| Class II | 34 | 15 |
| Class III | 27 | 11 |
| Toal amplified bands | 116 | 80 |
| Toal methylated bands | 61 | 26 |
| Fully methylated ratio (%) | 23.28 | 13.75 |
| MSAP ratio (%) | 52.59 | 32.50 |

Note: I belts were present in both *EcoRI/Hpa II* and *EcoRI/Msp I* digestion combination. II belts were present only in *EcoRI/Hpa II*. III belts were present only in *EcoRI/Msp I*. Total amplified bands = I + II + III, Total methylated bands = II + III, Fully methylated ratio (%) = III / (I + II + III) \* 100%, MSAP (%) = (II + III) / (I + II + III) \* 100%

### Effects of 5-azaC on the morphology, methylation level, and iridoid glycosides content of R. glutinosa roots

To further investigate the impact of DNA methylation on the accumulation of iridoid glycosides, 50 μМ 5-azaC was utilized to treat the roots of *R. glutinosa* from Wenxian during the I stage. The application of 5-azaC resulted in a significant increase in both root length and maximum cross-sectional diameter compared to the control group (Figure 4A-C). Both the fully methylated ratio and the MSAP ratio exhibited a marked decrease under 5-azaC treatment relative to the control group (Figure 4D, Figure S11). Analysis of iridoid glycoside levels indicated a substantial increase in iridoid glycoside content following exposure to 5-azaC compared to the control group (Figure 4E). It is postulated that the genomic demethylation induced by 5-azaC enhances the expression of genes involved in the biosynthesis of iridoid glycosides, thereby promoting their accumulation.

**Figure 4.**
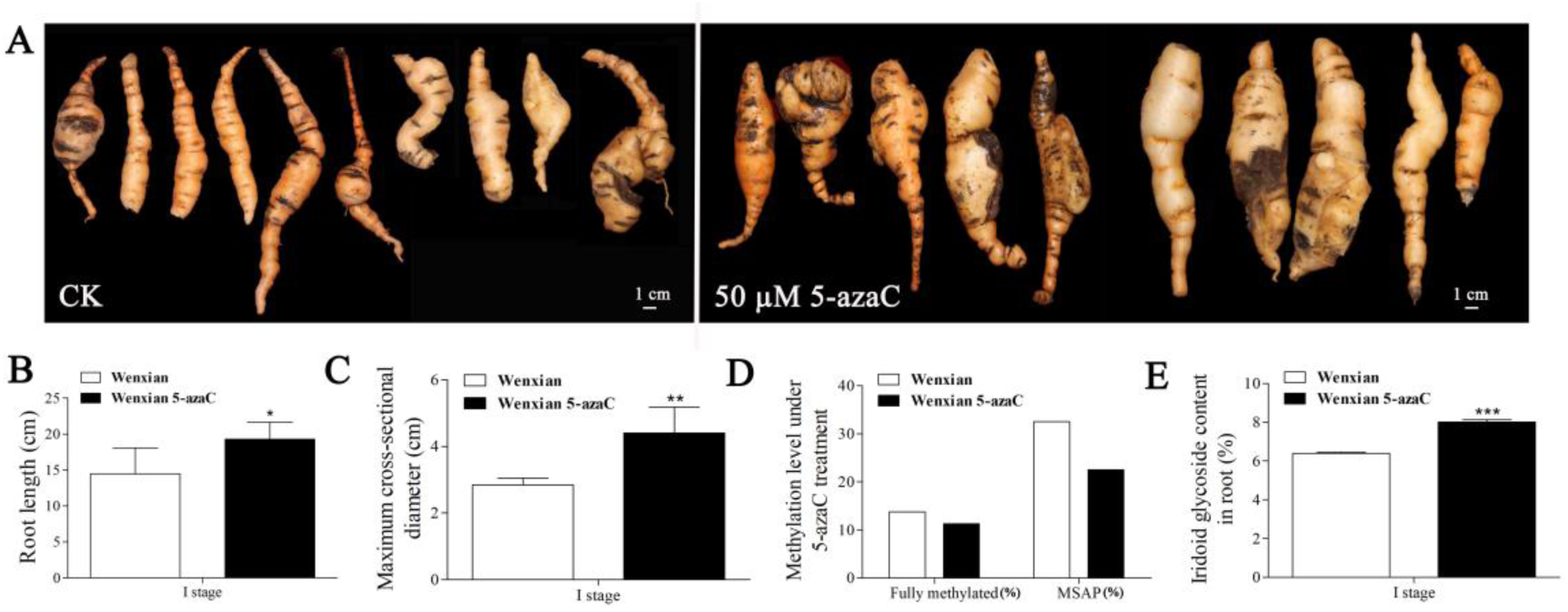
Morphology, methylation level, and iridoid glycosides content of *R. glutinosa* roots at E stage in Wenxian under 5-azaC treatment. (A-C) Morphological characteristics of *R. glutinosa*roots. (D) Methylation level of *R. glutinosa* roots. (E) The content of iridoid glycosides. The data are shown as the mean ± SD from three independent biological replicates (*, *P* < 0.05; **, *P* < 0.01; ***, *P* < 0.001; Student’s *t*-test).

### DEGs involved in the biosynthesis of iridoid glycosides under 5-azaC treatment

RNA-seq was utilized to perform a thorough investigation into the effects of DNA demethylation on the transcriptional activity of genes involved in the biosynthesis of iridoid glycosides. A total of 6,551 DEGs were identified following treatment with 50 μM 5-azaC, which included 5,248 upregulated genes and 1,313 downregulated genes (Figure S12A). GO term analysis indicated significant enrichment in categories such as metabolic process, cellular process, and biological regulation within the biological process category. The cellular component category showed enrichment in terms such as cell, organelle, and membrane, while the molecular function category highlighted enriched components including binding, catalytic activity, and transporter activity (Figure S12B). KEGG analysis indicated significant enrichment in pathways such as plant hormone signal transduction, MAPK signaling pathway, plant-pathogen interaction, starch and sucrose metabolism, and ubiquitin-mediated proteolysis (Figure S12C).

In the biosynthetic pathway of iridoid glycosides, 357 unigenes have been identified, of which 63 were up-regulated and 12 were down-regulated (Table S7). The cluster heatmap revealed that 5-azaC treatment led to the up-regulation of 4 *DXS*, 1 *DXR*, 1 *HDR*, and 3 *GPPS* associated with the MEP pathway. Downstream in the biosynthetic pathway, 28 unigenes were up-regulated, while only 5 unigenes related to *CPR*, *7-DLGT*, *7-DLH*, and *ALDH* were down-regulated (Figure 5A). UGTs play a crucial role in the glycosylation process, which is the final step in terpene biosynthesis. A total of 109 *UGTs* were identified, with 23 being up-regulated and 7 down-regulated (Figure 5B). Additionally, the TPS gene family, which is essential for generating diverse terpene structures, comprised 57 members, with 3 exhibiting up-regulation (Figure 5C). Furthermore, the expression levels of the methyltransferase genes were significantly decreased under 5-azaC treatment (Figure S12D).

**Figure 5.**
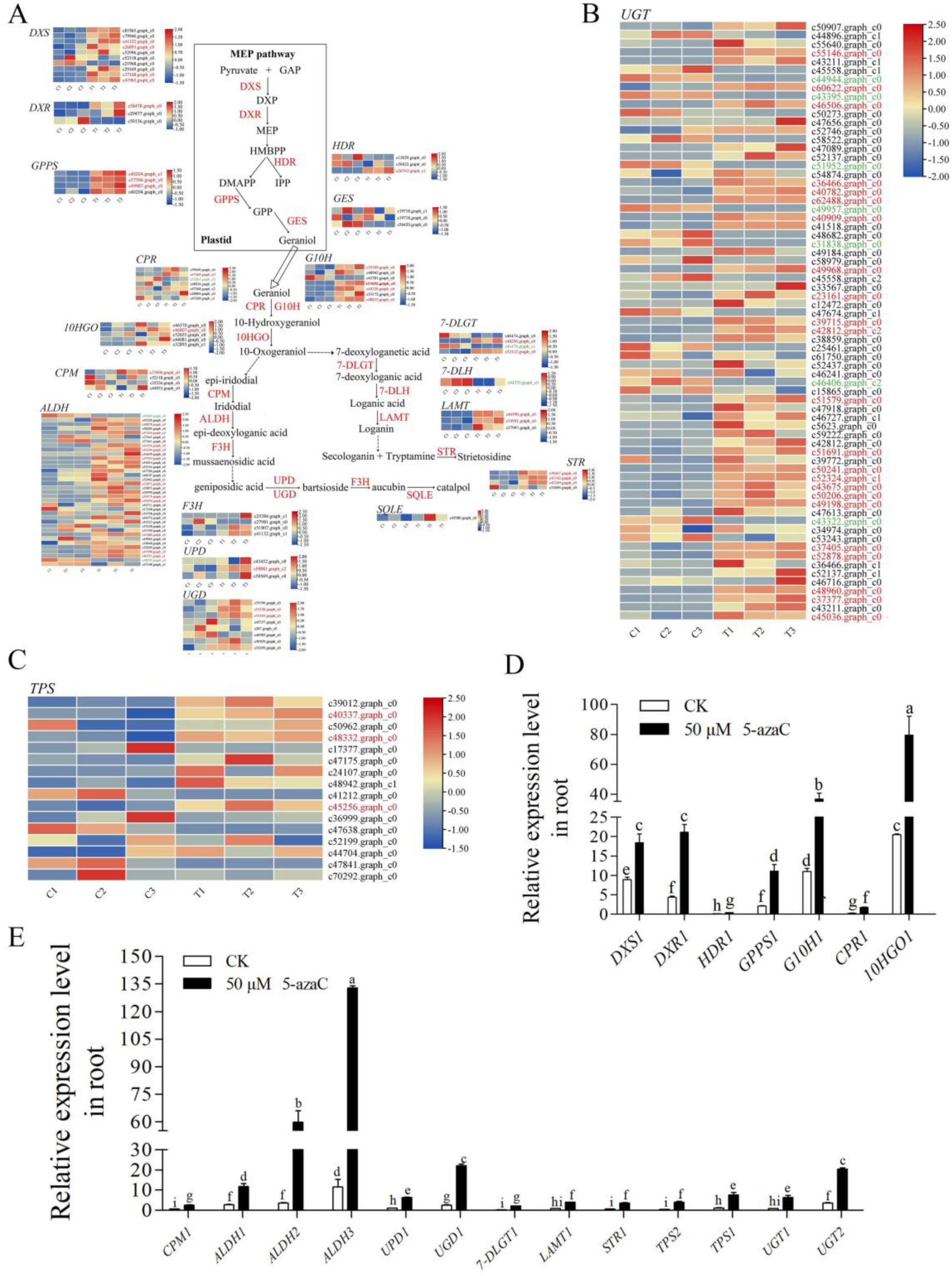
The unigenes relevant to iridoid glycoside in *R. glutinosa*. (A), (B) and (C) The expression heatmap of iridoid glycoside relevant unigenes. Blue and red color refers to the down or up expression of unigenes Each line represents the expression of each unigene in different groups and the expression values in row scale were normalized. Red unigene id indicates up-regulated, green unigene id means down-regulated and black unigene id indicates normal regulation. (D) and (E) qRT-PCR of unigenes relevant to iridoid glycoside under 5-azaC treatment. The *UPD1* expression level in the control group set to 100%. The error bar is standard error of mean, and the lower letter above the bar indicates the significant difference (*P* < 0.05).

To validate the RNA-seq results, significantly upregulated unigenes following 5-azaC treatment were confirmed using qRT-PCR. Most unigenes exhibited a 2- to 10-fold increase in expression, with *G10H1*, *10HGO1*, *ALDH2*, and *ALDH3* showing the most significant induction (Figure 5D, E).

### Differentially expressed TFs related to iridoid glycosides biosynthetic

TFs play a crucial role in regulating the expression of downstream enzyme genes involved in iridoid glycoside biosynthesis. A total of 1,215 unigenes were identified as TFs and categorized into 27 distinct families, with 139 exhibiting significant upregulation and 32 showing downregulation (Figure 6A, B; Table S8). qRT-PCR validation revealed that the expression levels of TFs were notably enhanced by 5-azaC, with AP2/ERF2, bHLH1, bZIP3, C2H21, HSF3, MYB4, and WRKY3 exhibiting the highest expression values within their respective gene families (Figure 6C, D). Furthermore, an interaction network involving 114 TFs and 28 enzymes was constructed. Ten TF proteins from the E2F/DP, bHLH, HSF, MYB, C3H, bZIP, and C2H2 families were predicted to interact with the enzymes ALDH13, HDR1, G10H4, DXR1, G10H3, and UPD1. Notably, DXR1 and HDR1, which are key enzymes in the upstream pathway of iridoid glycoside biosynthesis, were regulated by C3H1, E2F/DP1, and MYB1/2, respectively. G10H serves as the essential rate-limiting enzyme in monoterpene biosynthesis, with G10H3 and G10H4 being regulated by HSF2, C2H2_2, bZIP3, C3H1, and MYB2. ALDH13, which catalyzes the conversion of iridodial to epi-deoxyloganic acid, was regulated by C3H1 and HSF1/2/3 (Figure 6E).

**Figure 6.**
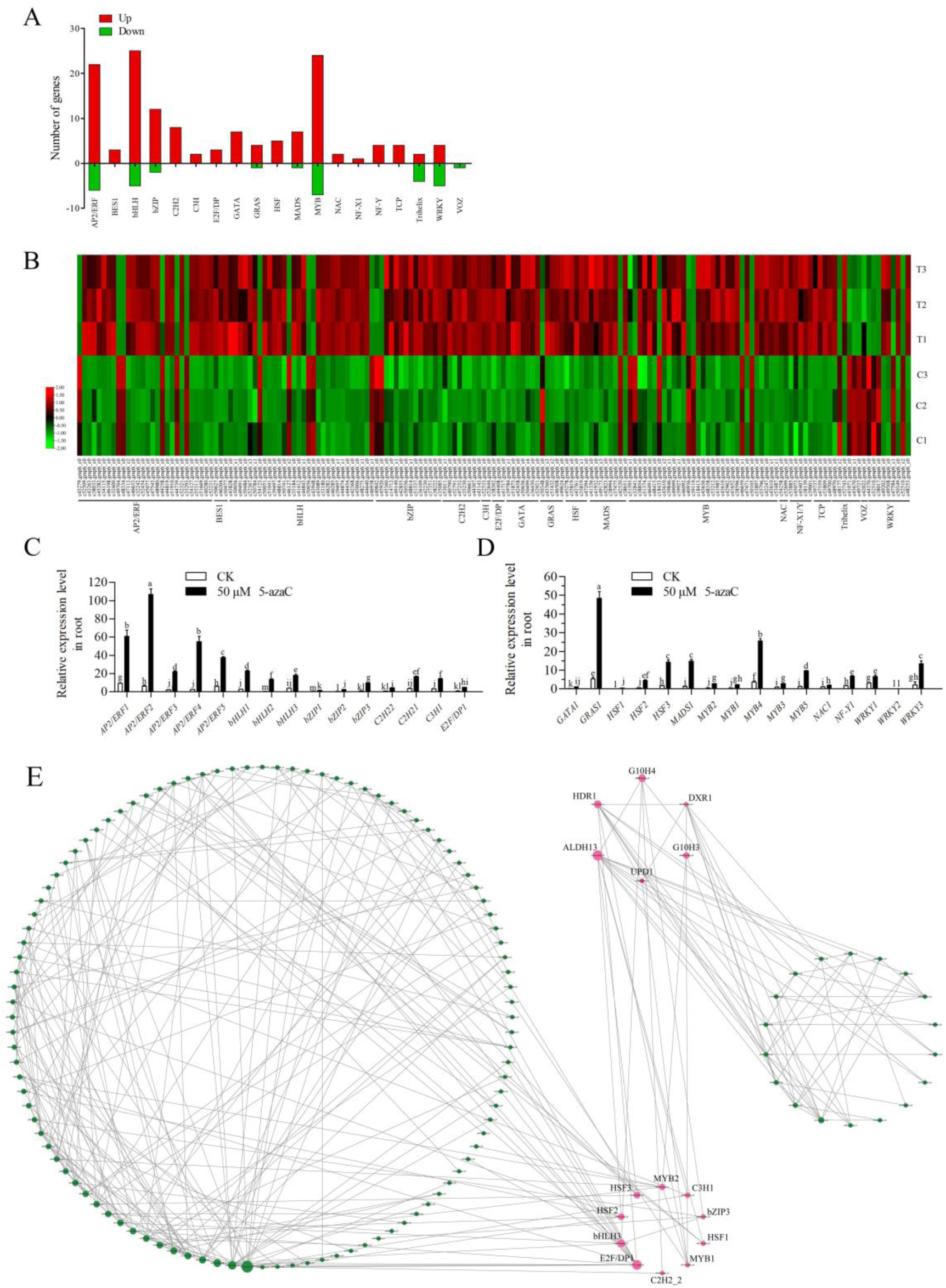
The DEGs of TFs in *R. glutinosa*. (A) The number of up- and down-regulation of TFs. Red color presents up regulation unigenes and green color indicates down regulation; (B) The expression heatmap of TFs. Red and green color refers to the up or down expression of TFs, respectively. Each line represents the expression of each unigene in different groups and the expression values in row scale were normalized. (C) and (D) qRT-PCR of TFs under 5-azaC treatment. The error bar is standard error of mean, and the lower letter above the bar indicates the significant difference (P < 0.05). (E) The PPI analysis of TFs and iridoid glycoside relevant unigenes. Pink color presents directly interaction relationship, and green color presents mediately interaction relationship. The size of the circle indicates the number of connections.

### Differential expression proteins under 5-azaC treatment

PPI analysis indicate that 5-azaC induces the upregulation of *DXR1*, *HDR1*, *G10H3*/*4*, *UPD1*, and *ALDH13*, suggesting potential co-regulation by TFs during the biosynthesis of iridoid glycosides. Furthermore, the study delves into the effect of 5-azaC on the biosynthesis of iridoid glycosides using iTRAQ (isobaric tags for relative and absolute quantitation). A total of 69 differentially expressed proteins (DEPs) were identified, meeting the criteria of a fold change (FC) greater than 1.2 or less than 0.83, along with statistical significance (*P* < 0.05). Among these, 43 proteins were upregulated, while 26 were downregulated following 5-azaC treatment (Table S9). Specifically, two proteins associated with the G10H and ALDH gene families, which are critical in the iridoid glycoside biosynthetic pathway, were detected. Integration analysis confirmed that these G10H and ALDH proteins correspond to *RgG10H4* (c55692.graph_c0) and *RgALDH74* (c48592.graph_c0), respectively, as identified in the transcriptome.

### Characteristics, expression, and subcellular localization of RgG10H4

Geraniol serves as the initiator of the iridoid glycosides biosynthesis pathway and undergoes hydroxylation, catalyzed by the first rate-limiting enzyme, G10H, to form 10-hydroxygeraniol. The CDS of *RgG10H4* spans 1,473 bp, translating into a protein composed of 491 amino acids. This sequence is characterized by the presence of a signal peptide and a transmembrane domain. The expression level of *RgG10H4* peaked at I stage in roots, followed by a significant decrease in expression at M stage (Figure 7A). Subcellular localization revealed that the GFP signal from the control was distributed throughout the entire cell, whereas the RgG10H4-GFP signal was specifically localized to the endoplasmic reticulum (ER) (Figure 7B).

**Figure 7.**
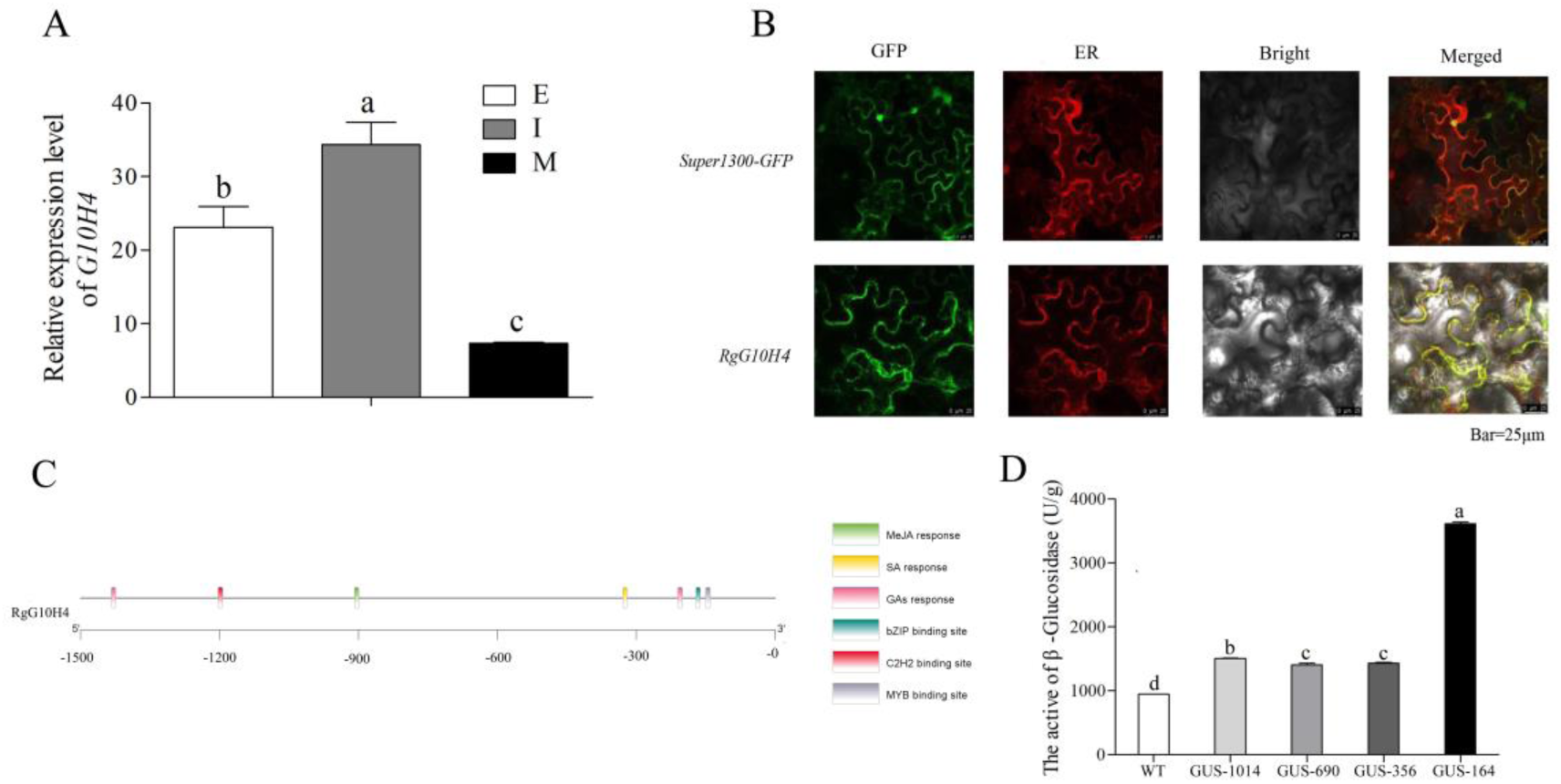
Analysis of *RgG10H4* in *R. glutinosa*. (A) Expression pattern of *RgG10H4*. The data are shown as the mean ± SD from three independent biological replicates. The lower letter above the bar indicates the significant difference (*P* < 0.05). (B) Subcellular localization of RgG10H4. The yellow region indicates ER localization. GFP: Green fluorescent protein. Bar = 25 μm. (C) Analysis of *cis*-acting elements in *RgG10H4* promoter. (D) Analysis of GUS activity in the *RgG10H4* promoter. The data are shown as the mean ± SD from three independent biological replicates. The lower letter above the bar indicates the significant difference (*P* < 0.05).

### Promoter cloning, cis-acting element, and GUS activity of RgG10H4

The fusion primer and nested integrated PCR (FPNI-PCR) were employed to clone the promoter of *RgG10H4*, resulting in the acquisition of a 1,676 bp promoter sequence after three rounds of FPNI-PCR (Figure S13A, B). An examination of the *cis*-acting elements within the *RgG10H4* promoter region revealed the presence of responsive elements for MeJA, GAs, and SA, as well as binding sites for MYB, C2H2, and bZIP TFs (Figure 7C). The *RgG10H4* promoter was fragmented into segments of 1,014 bp, 690 bp, 356 bp, and 164 bp, which were subsequently ligated with the pCAMBIA 1301 vector to create GUS fusion constructs. Following transient transfection of tobacco leaves, β-glucosidase activity was measured, demonstrating maximal enzyme activity in the -164 bp truncated fragment. These findings indicate that the critical functional domain of the *RgG10H4* promoter is located within the proximal 164 bp upstream region (Figure 7D).

### RgMYB2 binds to the promoter regions of RgG10H4

Molecular docking was performed to investigate potential binding sites between the RgMYB2, RgC2H2-2, and RgbZIP3 proteins and the promoter region of *RgG10H4*. The confidence scores for the interactions between the *RgG10H4* promoter and the proteins RgMYB2, RgC2H2-2, and RgbZIP3 were calculated to be 0.8993, 0.8935, and 0.8558, respectively. Further analysis revealed the formation of stable complexes between the promoter region of *RgG10H4* and the domains of RgMYB2, RgC2H2-2, and RgbZIP3, facilitated by strong hydrogen bond interactions (Figure 8A). The yeast one-hybrid (Y1H) assay confirmed that the RgMYB2 protein specifically bound to the fragment that containing the TAACCA motif of the *RgG10H4* promoter on the SD/-Leu+AbA (200 ng/ml) medium (Figure 8B). These results substantiate the direct interaction between RgMYB2 and the promoter sequences of *RgG10H4*.

**Figure 8.**
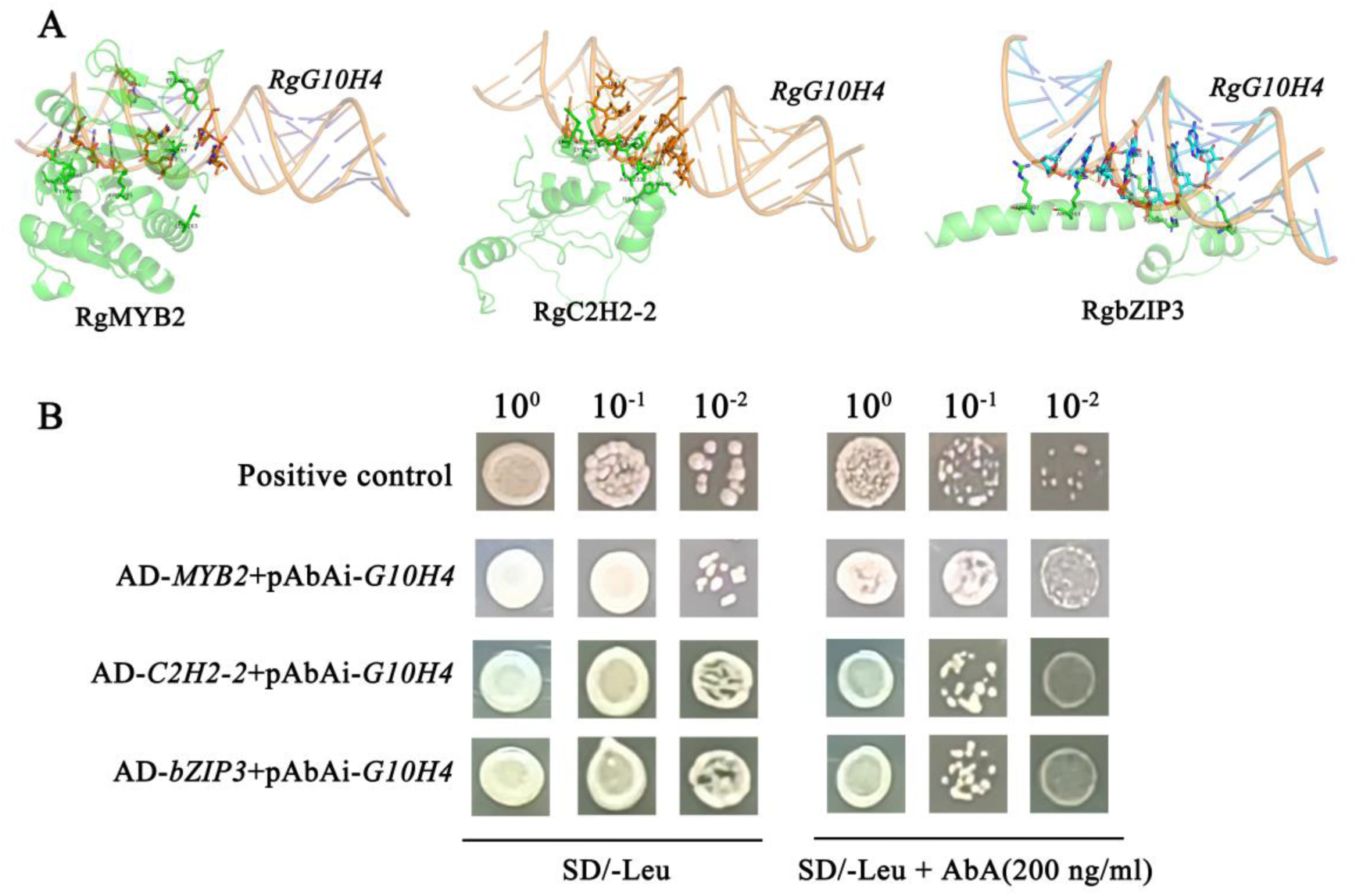
The RgMYB2 binds to the promoter regions of *RgG10H4*. (A) The molecular docking 3D structure of the interactions between RgMYB2, RgC2H2-2, RgbZIP3 proteins and *RgG10H4* promoter. The yellow dashed line represents hydrogen bonding interactions. (B) The Y1H experiment was performed to investigate the interaction of *RgMYB2*, *RgC2H2-2*, and *RgbZIP3* with the promoter region of *RgG10H4*. The promoter region of *RgG10H4* was cloned into the pAbAi vector, while the CDS of *RgMYB2*, *RgC2H2-2*, and *RgbZIP3* were inserted into the pGADT7 vector for the assay. Yeast cells were diluted with distilled water (10^0^ to 10^-2^) and cultured on SD/-Leu medium supplemented with 200 ng/mL of Aureobasidin A (AbA).

## Discussion

PSMs are a class of non-essential, small molecular organic compounds produced through secondary metabolism, which are crucial for the normal functioning of cellular activities, as well as for plant growth and development (Chiocchio et al., 2021). The accumulation of secondary metabolites in plants, which serves as an important material basis, is influenced by environmental factors and phenological stages (Yang et al., 2018; Kumari et al., 2024; Liu et al., 2024). Here, we found that at the same growth stage, the morphology of *R. glutinosa* roots from Wenxian was superior to that from Xinxiang, and the content of iridoid glycosides was also higher in the Wenxian samples compared to those from Xinxiang (Figure 1). Metabolomic analyses identified 13 types of iridoid compounds in *R. glutinosa* roots, with 9 of these compounds occurring at higher concentrations in the Wenxian production area. Furthermore, the expression levels of 43 key genes involved in the biosynthesis of iridoid glycosides were elevated in the roots of *R. glutinosa* from the Wenxian region (Figure 3C). Previous studies have shown that the level of DNA methylation in plants varies across different environments, representing a crucial factor that influences the accumulation of PSMs (Xu et al., 2020; Zhao et al., 2023). We suggest that the higher contents of iridoid glycosides in Wenxian may be attributed to differences in DNA methylation levels caused by varying growth environments.

In plants, variations in DNA methylation levels influence growth, development, and the accumulation of PSMs (Zhang et al., 2018). In *Arabidopsis thaliana*, treatment with 5-azaC promoted the formation of adventitious roots in mature plants (Massoumi et al., 2017). The application of 5-azaC significantly enhanced the stem regeneration efficiency in soybean genotypes with low regeneration capacity (Zhao et al., 2019). However, excessive concentrations of 5-azaC markedly inhibited the height of *Pogostemon cablin*, leading to abnormalities in leaf and corolla development (Luo et al., 2021). In the study, we found that the MSAP% of *R. glutinosa* root in Wenxian was significantly lower than that in Xinxiang (Table 1). Treatment with 5-azaC enhanced roots development, reduced DNA methylation levels, and significantly increased the content of iridoid glycosides in *R. glutinosa* roots from Wenxian (Figure 4). Similarly, the DNA methylation levels in birch were significantly reduced, while the triterpenoid content was significantly increased following treatment with 5-azaC (Zeng et al., 2020).

Currently, transcriptomics is employed to identify pivotal genes involved in the biosynthetic pathways of iridoids. In *Gardenia jasminoides*, fifteen enzyme genes associated with iridoid synthesis were identified, including DXS, DXR, G10H, 10HGO, and GPPS, among others. The expression levels of these genes exhibited a positive correlation with fruit development (Pan et al., 2021). A total of 23 key enzyme-encoding genes involved in iridoid biosynthesis were identified in *Phlomoides rotata*, with the expression levels of *HDR2*, *10HGO1*, *GPPS2*, and *G10H1* being notably higher. Further analysis revealed that *DXS1*, *IDI1*, *10HGO1*, and *G10H2* were significantly correlated with the accumulation of iridoids (Wang et al., 2024). Moreover, 1,395 TFS were identified in *Gentiana macrophylla*, among which *GmbHLH20* and *GmMYB5* may positively regulate the expression of *GmDXS*, *GmHDR1*, *GmG10H*, *Gm10HGO*, *GmGPPS*, and *GmIS3* (Fu et al., 2024). Herein, the key genes involved in the iridoid glycoside pathway were identified in *R. glutinosa* using RNA-seq. A total of 357 genes related to iridoid glycoside synthesis pathway were identified, of which 63 genes were upregulated following 5-azaC treatment. Notably, four genes encoding G10H and one gene encoding 10HGO, which are key enzymes linking the upstream MEP pathway to downstream monoterpene biosynthesis, were significantly upregulated (Figure 5A-C). Subsequently, the genes with higher expression levels were selected for qRT-PCR validation, and the results were consistent with the transcriptome analysis (Figure 5D,E). Similarly, the expression levels of genes related to flavonoid and anthocyanin metabolic pathways were significantly elevated in apples following treatment with 5-azaC, including flavonol synthases (*FLSs*), leucoanthocyanidin reductases (*LARs*), and anthocyanidin reductases (*ANRs*). This upregulation likely contributed to a marked enhancement in flavonoid accumulation (Ma et al., 2018). Another study revealed that the expression levels of flavonoid synthetase genes, including *CHSs*, *PALs*, *F2Hs*, and *PTs* were induced by 5-azaC in *Glycyrrhiza inflata*. Moreover, the expression levels of TFs, such as MYB, WRKY, ERF, NAC, and bZIP, were significantly increased, resulting in enhanced flavonoid accumulation (Ma et al., 2024). In *Salvia miltiorrhiza*, elevated expression of *SmCMT2*, *SmDDM1*, *SmAGO4*, and *SmDRM1* resulted in increased genomic DNA methylation levels. High methylation was associated with decreased expression of numerous genes involved in the biosynthesis of tanshinones and salvianolic acids, including *SmCPS5*, *SmCYP71*, *SmGGPPS1*, *SmGPPS*, *SmHPPR*, and *SmHPPD*, ultimately leading to reduced accumulation of tanshinones and phenolic acids (He et al., 2024). Interestingly, the application of 5-azaC to treat the hairy roots of *Salvia miltiorrhiza* significantly increased the contents of cryptotanshinone, tanshinone I, dihydrotanshinone, and tanshinone IIA. Further analysis revealed that the DNA methylation levels of key enzyme genes in the tanshinone biosynthesis pathway, including *DXS*, *DXR*, *GGPPS*, *CPS*, *HDR*, and *CYP76*, were downregulated, while their expression levels were upregulated (Li et al., 2023). These findings suggest that DNA demethylation induced by 5-azaC treatment enhances the expression of key enzyme genes involved in the secondary metabolic pathway, resulting in an accumulation of PSMs.

TFs facilitate the coordinated expression of multiple genes within the secondary metabolite synthesis pathway, thereby initiating the pathway and enhancing the accumulation of PSMs (Zha et al., 2017; Sanchez-Muñoz et al., 2018; Wei et al., 2022; Liu et al., 2024). In *Dendrobium officinale*, *DoWRKY*, *DoMYB*, *DobHLH*, *DobZIP*, and *DoERF* established a regulatory network with *DoTPS04*, *DoTPS10*, and *DoTPS07* to modulate terpenoid biosynthesis (Li et al., 2021). We found that 161 TFs were regulated following treatment with 5-azaC. Of these, 114 TF proteins were mapped using the STRING database (Figure 6). PPI analysis indicated that these TFs, including E2F/DP, bHLH, HSF, MYB, C3H, bZIP, and C2H2, may interact with key enzymes involved in the biosynthesis of iridoid glycosides (Figure 6E). Subsequently, proteomic analysis revealed that while the expression of numerous key enzyme genes involved in the iridoid glycoside synthesis pathway was enhanced by 5-azaC, only 43 proteins exhibited a significant increase in abundance. Notably, RgG10H4 and RgALDH74 showed particularly pronounced increases. Geranyl 10-hydroxylase (G10H) is a cytochrome P450 monooxygenase that acts as the initial rate-limiting enzyme in the synthesis of terpenoid skeletons through the MEP pathway, thereby playing a vital role in the production of iridoid glycosides (Collu et al., 2001; Pu et al., 2021). In *Catharanthus roseus*, the promoter of *G10H* is abundant in binding sites for MYB, WRKY, and Dof TFs. GUS assays demonstrated that the active regions of the promoter are primarily located in the upstream region of the transcription start site (TSS) at -103 bp and within the -318 to -147 bp interval (Suttipanta et al., 2007). We found that the *RgG10H4* promoter contains a high density of hormone regulatory elements and transcription factor binding sites. The most active region is located at -164 bp upstream, where the core MYB sequence (TAACCA) is precisely located (Figure 7C, D).

Furthermore, ample studies have indicated that TFs regulate the expression of key enzyme genes by binding to specific sequences in their promoters, thereby influencing the accumulation of PSMs (Zhao et al., 2023). AaMYC2 and AabZIP1 serve as positive regulators of the artemisinin biosynthesis pathway by enhancing the expression of *AaWRKY1*. In turn, the AaWRKY1 protein directly binds to the W boxes in the promoters of *AaCYP71AV1* and *AaORA*, further facilitating the accumulation of artemisinin in *Artemisia annua* (Chen et al., 2017). In *S. miliorrhiza*, *SmERF128* and *SmERF152* regulate the biosynthetic pathway of tanshinone. The SmERF128 protein binds to the promoter-specific elements of *SmCPS1*, *SmKSL1*, and *SmCYP76AH1*, thereby activating their expression and promoting the accumulation of tanshinone (Ji et al., 2016; Zhang et al., 2019). MYB TFs play a crucial role in regulating the biosynthetic pathways of monoterpenes, sesquiterpenes, and triterpene saponins. Research has shown that *AaMYB1* positively influences artemisinin biosynthesis, while *AaMYB15* functions as a negative regulator. Specifically, *AaMYB15* reduces transcriptional activity by binding to the promoters of *ADS*, *CYP*, *DBR2*, *ALDH1*, and *AaORA*, thereby inhibiting artemisinin synthesis (Wu et al., 2021). In a separate study, *PgMYB2* was found to play a positive regulatory role in the synthesis of ginsenosides in *Panax ginseng*. PgMYB2 enhances the expression of *PgDDS* by binding to its promoter, resulting in increased ginsenoside accumulation (Liu et al., 2019). In the study, the RgMYB2 protein specifically binds to the TAACCA motif in the *RgG10H4* promoter, forming a stable hydrogen bond structure (Figure 8). Overall, treatment with 5-azaC enhanced the expression of both *RgMYB2* and *RgG10H4*, suggesting that the *RgMYB2*-*RgG10H4* module may play a positive role in the biosynthesis of iridoid glycosides.

In conclusion, our study is the first to elucidate the role of DNA methylation in the accumulation of iridoid glycosides in *R. glutinosa* by integrating transcriptomics, proteomics, and metabolomics. The findings demonstrate that appropriate DNA demethylation enhances the growth and development of *R. glutinosa* roots and stimulates the expression of key enzyme genes and TFs involved in the iridoid glycoside biosynthesis pathway, leading to an increased accumulation of iridoid glycosides. Further analysis suggests that the *RgMYB2*-*RgG10H4* module plays a positive regulatory role in the biosynthesis of iridoid glycosides.

## Acknowledgment

The study was supported by “The High-Performance Computing Center of Henan Normal University”.

## Supporting Information

### Supplemental Figures

**Supplemental Figure S1.** Content and proportions of secondary metabolites in *R. glutinosa* roots.

**Supplemental Figure S2.** Principal component analysis of secondary metabolites in *R. glutinosa* roots from the Wenxian and Xinxiang production areas.

**Supplemental Figure S3.** Hierarchical clustering analysis of secondary metabolites in *R. glutinosa* roots from the Wenxian and Xinxiang production areas.

**Supplemental Figure S4.** Correlation heatmap analysis of secondary metabolites in *R. glutinosa* roots from the Wenxian and Xinxiang production areas.

**Supplemental Figure S5.** A volcanic map illustrating DAMs in *R. glutinosa* roots from the Wenxian and Xinxiang production areas.

**Supplemental Figure S6.** Hierarchical clustering analysis of DAMs in *R. glutinosa* roots from the Wenxian and Xinxiang production areas.

**Supplemental Figure S7.** A volcanic map illustrating DEGs in *R. glutinosa* roots from the Wenxian and Xinxiang production areas.

**Supplemental Figure S8.** Clustering heatmap analysis of DEGs in *R. glutinosa* roots from the Wenxian and Xinxiang production areas.

**Supplemental Figure S9.** Genome methylation ratio in *R. glutinosa* roots from the Xinxiang and Wenxian production areas.

**Supplemental Figure S10.** Clustering heatmap analysis of methyltransferase and demethyltransferase genes in the roots of *R. glutinosa* from the Wenxian and Xinxiang production areas.

**Supplemental Figure S11.** Genome methylation ratio in *R. glutinosa* roots from Wenxian production areas under 5-azaC treatment.

**Supplemental Figure S12.** Analysis of DEGs in *R. glutinosa* under 5-azaC treatment.

**Supplemental Figure S13** The promoter of *RgG10H4* was cloned using FPNI-PCR techniques. The 1676 bp promoter sequence was obtained through three rounds of amplification.

### Supplemental Tables

**Supplemental Table 1** The primers used in this study.

**Supplemental Table 2** Type and relative abundance of all metabolites in *R. glutinosa* roots.

**Supplemental Table 3** List of DAMs in *R. glutinosa* roots.

**Supplemental Table 4** List of iridoid glycoside in *R. glutinosa* roots.

**Supplemental Table 5** DEGs of iridoid glycoside biosynthesis in *R. glutinosa* roots.

**Supplemental Table 6** PPI of unigenes involved in iridoid glycoside biosynthesis.

**Supplemental Table 7** DEGs involved in iridoid glycoside biosynthesis under 5-azaC treatment.

**Supplemental Table 8** DEGs of TFs under 5-azaC treatment.

**Supplemental Table 9** DEPs in *R. glutinosa* roots under 5-azaC.

## Author Contributions

Tianyu Dong: conceptualization; Tianyu Dong, and Yajie Du: methodology; Tianyu Dong, and Yajie Du: formal analysis; Tianyu Dong, and Jie Guo: investigation; Qingxiang Yang, Jingjing Xing, and Hongying Duan: resources; Tianyu Dong, and Tingting Huang: data curation; Tianyu Dong: writing - original draft; Jiuchang Su, Qingxiang Yang, and Peilei Chen: writing - review & editing; Tianyu Dong: visualization; Jingjing Xing, and Hongying Duan: supervision; Jingjing Xing, and Hongying Duan: funding acquisition

## Data Available

All of the relevant data and figures in this study can be found in the article and its supplementary data.

## Abbreviations

5-azaC: 5-Azacytidine
PSMs: Plant secondary metabolites
HCA: hierarchical clustering analysis
TFs: transcription factors
DEGs: differentially expressed genes
DAMs: differentially abundant metabolites
MSAP: methylation-sensitive amplified polymorphism
FPNI-PCR: fusion primer and nested integrated PCR
β-GC: β-glucosidase
CS: confidence score
AbA: Aureobasidin A
G10H: geranyl 10-hydroxylase
TSS: transcription start site
Y1H: yeast one-hybrid

## REFERENCES

1. Chang Y, Zhu C, Jiang J, Zhang H, Zhu J, Duan C. 2020. Epigenetic regulation in plant abiotic stress responses. J. Integr. Plant Biol. 62, 563–580.

2. Chen M, Yan T, Shen Q, et al. 2017. GLANDULAR TRICHOME-SPECIFIC WRKY 1 promotes artemisinin biosynthesis in *Artemisia annua*. New Phytol. 214, 304–316.

3. Chen P, Wei X, Qi Q, Jia W, Zhao M, Wang H, Zhou Y, Duan H. 2021. Study of terpenoid synthesis and prenyltransferase in roots of *Rehmannia glutinosa* based on iTRAQ quantitative proteomics. Front. Plant Sci. 12, 693758.

4. Chen S, Zhou Y, Chen Y, Gu J. 2018. Fastp: an ultra-fast all-in-one FASTQ preprocessor. Bioinformatics 34, i884–i890.

5. Chen W, Gong L, Guo Z, Wang W, Zhang H, Liu X, Yu S, Xiong L, Luo J. 2013. A novel integrated method for large-scale detection, identification, and quantification of widely targeted metabolites: application in the study of rice metabolomics. Mol. Plant 6, 1769–1780.

6. Chen Y, Li D, Xu Y, Lu Z, Luo Z. 2024. 5-Azacytidine accelerates mandarin fruit post-ripening and enhances lignin-based pathogen defense through remarkable gene expression activation. Food Chem. 458, 140261.

7. Chiocchio I, Mandrone M, Tomasi P, Marincich L, Poli F. 2021. Plant secondary metabolites: an opportunity for circular economy. Molecules 26, 495.

8. Collu G, Unver N, Peltenburg-Looman AM, van der Heijden R, Verpoorte R, Memelink J. 2001. Geraniol 10-hydroxylase, a cytochrome P450 enzyme involved in terpenoid indole alkaloid biosynthesis. FEBS Lett. 508, 215–220.

9. Dong T, Song S, Wang Y, Yang R, Chen P, Su J, Ding X, Liu Y, Duan H. 2022. Effects of 5-azaC on iridoid glycoside accumulation and DNA methylation in *Rehmannia glutinosa*. Front. Plant Sci. 13, 913717.

10. Faria DV, de Freitas Correia LN, Batista DS, Vital CE, Heringer AS, De-la-Peña C, Cardoso-Costa MG, Guerra MP, Otoni WC. 2020. 5-Azacytidine downregulates the *SABATH* methyltransferase genes and augments bixin content in *Bixa orellana* L. leaves. Plant Cell Tiss. Organ. Cult. 142, 425–434.

11. Ferrandino A, Pagliarani C, Pérez-Álvarez EP. 2023. Secondary metabolites in grapevine: crosstalk of transcriptional, metabolic and hormonal signals controlling stress defence responses in berries and vegetative organs. Front. Plant Sci. 14, 1124298.

12. Fraga CG, Clowers BH, Moore RJ, Zink EM. 2010. Signature-discovery approach for sample matching of a nerve-agent precursor using liquid chromatography-mass spectrometry, XCMS, and chemometrics. Anal. Chem. 82, 4165–4173.

13. Fu H, Wang Y, Mi F, Wang L, Yang Y, Wang F, Yue Z, He Y. 2024. Transcriptome and metabolome analysis reveals mechanism of light intensity modulating iridoid biosynthesis in *Gentiana macrophylla* Pall. BMC Plant Biol. 24, 526.

14. Grabherr M, Haas B, Yassour M, et al. 2011. Full-length transcriptome assembly from RNA-Seq data without a reference genome. Nat. Biotechnol. 29, 644–652.

15. Han M, Lin S, Zhu B, et al. 2024. Dynamic DNA methylation regulates season-dependent secondary metabolism in the new shoots of tea plants. J. Agric. Food Chem. 72, 3984–3997.

16. Hao M, Zhou Y, Zhou J, Zhang M, Yan K, Jiang S, Wang W, Peng X, Zhou S. 2020. Cold-induced ginsenosides accumulation is associated with the alteration in DNA methylation and relative gene expression in perennial American ginseng (*Panax quinquefolius* L.) along with its plant growth and development process. J. Ginseng Res. 44, 747–755.

17. Hartmann T. 2007. From waste products to ecochemicals: fifty years research of plant secondary metabolism. Phytochemistry 68, 2831–2846.

18. He X, Chen Y, Xia Y, et al. 2024. DNA methylation regulates biosynthesis of tanshinones and phenolic acids during growth of *Salvia miltiorrhiza*. Plant Physiol. 194, 2086–2100.

19. Jacek RW, Alexandre Z, Nagarjuna N, Matthias M. 2009. Universal sample preparation method for proteome analysis. Nat. Methods 6, 359–362.

20. Jamloki A, Bhattacharyya M, Nautiyal MC, Patni B. 2021. Elucidating the relevance of high temperature and elevated CO_2_ in plant secondary metabolites (PSMs) production. Heliyon 7, e07709.

21. Jan R, Asaf S, Numan M, Lubna, Kim KM. 2021. Plant secondary metabolite biosynthesis and transcriptional regulation in response to biotic and abiotic stress conditions. Agronomy 11, 968.

22. Ji A, Luo H, Xu Z, Zhang X, Zhu Y, Liao B, Yao H, Song J, Chen S. 2016. Genome-wide identification of the AP2/ERF gene family involved in active constituent biosynthesis in *Salvia miltiorrhiza*. Plant Genome 9, 1877020.

23. Jia H, Zhang Z, Sadeghnezhad E, Pang Q, Li S, Pervaiz T, Su Z, Dong T, Fang J, Jia H. 2020. Demethylation alters transcriptome profiling of buds and leaves in ’Kyoho’ grape. BMC Plant Biol. 20, 544.

24. Kumari S, Nazir F, Maheshwari C, Kaur H, Gupta R, Siddique KHM, Khan MIR. 2024. Plant hormones and secondary metabolites under environmental stresses: Enlightening defense molecules. Plant Physiol. Biochem. 206, 108238.

25. Li J, Li C, Deng Y, Wei H, Lu S. 2023. Characteristics of *Salvia miltiorrhiza* methylome and the regulatory mechanism of DNA methylation in tanshinone biosynthesis. Hortic. Res. 10, uhad114.

26. Li N, Dong Y, Lv M, Qian L, Sun X, Liu L, Cai Y, Fan H. 2021. Combined analysis of volatile terpenoid metabolism and transcriptome reveals transcription factors related to terpene synthase in two cultivars of *Dendrobium officinale* flowers. Front. Genet. 12, 661296.

27. Li Y, Kong D, Fu Y, Sussman MR, Wu H. 2020. The effect of developmental and environmental factors on secondary metabolites in medicinal plants. Plant Physiol. Bioch. 148, 80–89.

28. Li Y, Zhai X, Ma L, Zhao L, An N, Feng W, Huang L, Zheng X. 2024. Transcriptome analysis provides insights into catalpol biosynthesis in the medicinal plant *Rehmannia glutinosa* and the functional characterization of *RgGES* genes. Genes (Basel) 15, 155.

29. Liu F, Xi M, Liu T, Wu X, Ju L, Wang D. 2024. The central role of transcription factors in bridging biotic and abiotic stress responses for plants’ resilience. New Crops 1, 100005.

30. Liu T, Luo T, Guo X, Zou X, Zhou D, Afrin S, Li G, Zhang Y, Zhang R, Luo Z. 2019. *PgMYB2*, a MeJA-responsive transcription factor, positively regulates the dammarenediol synthase gene expression in *Panax Ginseng*. Int. J. Mol. Sci. 20, 2219.

31. Liu Y, Chen S, Pal S, Yu J, Zhou Y, Phan Tran LS, Xia X. 2024. The hormonal, metabolic, and environmental regulation of plant shoot branching. New Crops 1, 100028.

32. Luo K, Ou X, Deng W, Liu X, He M, Zhang H, Yan H. 2021. Effects of 5-azacytidine on DNA methylation and index components in patchouliol-type *Pogostemon cablin*. China J. Chinese Materia Medica. 46, 4117–4123.

33. Ma C, Liang B, Chang B, Liu L, Yan J, Yang Y, Zhao Z. 2018. Transcriptome profiling reveals transcriptional regulation by DNA methyltransferase inhibitor 5-Aza-2’-Deoxycytidine enhancing red pigmentation in bagged “Granny Smith” apples (*Malus domestica*). Int. J. Mol. Sci. 19, 3133.

34. Ma X, Jiang N, Fu J, Li Y, Zhou L, Yuan L, Wang Y, Li Y. 2024. A cytosine analogue 5-azacitidine improves the accumulation of licochalcone A in licorice *Glycyrrhiza inflata*. J. Plant Physiol. 292, 154145.

35. Martínez-Rivas FJ, Blanco-Portales R, Molina-Hidalgo FJ, Caballero JL, Perez de Souza L, Alseekh S, Fernie AR, Muñoz-Blanco J, Rodríguez-Franco A. 2022. Azacytidine arrests ripening in cultivated strawberry (Fragaria × ananassa) by repressing key genes and altering hormone contents. BMC Plant Biol. 22, 278.

36. Massoumi M, Krens FA, Visser RG, De Klerk GM. 2017. Azacytidine and miR156 promote rooting in adult but not in juvenile *Arabidopsis* tissues. J. Plant Physiol. 208, 52–60.

37. Pan Y, Zhao X, Wang Y, Tan J, Chen D. 2021. Metabolomics integrated with transcriptomics reveals the distribution of iridoid and crocin metabolic flux in *Gardenia jasminoides* Ellis. PLoS One 16, e0256802.

38. Pu X, Dong X, Li Q, Chen Z, Liu L. 2021. An update on the function and regulation of methylerythritol phosphate and mevalonate pathways and their evolutionary dynamics. J. Integr. Plant Biol. 63, 1211–1226.

39. Sanchez-Muñoz R, Bonfill M, Cusidó RM, Palazón J, Moyano E. 2018. Advances in the regulation of in vitro paclitaxel production: Methylation of a Y-patch promoter region alters BAPT gene expression in taxus cell cultures. Plant Cell Physiol. 59, 2255–2267.

40. Saslis-Lagoudakis CH, Savolainen V, Williamson EM, Forest F, Wagstaff SJ, Baral SR, Watson MF, Pendry CA, Hawkins JA. 2012. Phylogenies reveal predictive power of traditional medicine in bioprospecting. Proc. Natl. Acad. Sci. U S A. 109, 15835–15840.

41. Schymanski EL, Jeon J, Gulde R, Fenner K, Ruff M, Singer HP, Hollender J. 2014. Identifying small molecules via high resolution mass spectrometry: communicating confidence. Environ. Sci. Technol. 48, 2097–2098.

42. Suttipanta N, Pattanaik S, Gunjan S, Xie C, Littleton J, Yuan L. 2007. Promoter analysis of the *Catharanthus roseus* geraniol 10-hydroxylase gene involved in terpenoid indole alkaloid biosynthesis. Biochim. Biophys. Acta. 1769, 139–148.

43. Thiebaut F, Hemerly AS, Ferreira PCG. 2019. A role for epigenetic regulation in the adaptation and stress responses of non-model plants. Front. Plant Sci. 10, 246.

44. Wang C, Tang L, Li L, Zhou Q, Li Y, Li J, Wang Y. 2020. Geographic authentication of *Eucommia ulmoides* leaves using multivariate analysis and preliminary study on the compositional response to environment. Front. Plant Sci. 11, 79.

45. Wang H, Zhu Y, Yuan P, Song S, Dong T, Chen P, Duan Z, Jiang L, Lu L, Duan H. 2021. Response of wheat DREB transcription factor to osmotic stress based on DNA methylation. Int. J. Mol. Sci. 22, 7670.

46. Wang L, Geng G, Xie H, Zhou L, He Y, Li Z, Qiao F. 2024. A transcriptomic and metabolomic study on the biosynthesis of iridoids in *Phlomoides rotata* from the Qinghai-Tibet Plateau. Plants (Basel) 13, 1627.

47. Wang S, Alseekh S, Fernie A, Luo J. 2019. The structure and function of major plant metabolite modifications. Mol. Plant 12, 899–919.

48. Wang S, Zhao X, Li C, Dong J, Ma J, Long Y, Xing Z. 2024. DNA methylation regulates the secondary metabolism of saponins to improve the adaptability of *Eleutherococcus senticosus* during drought stress. BMC Genomics 25, 330.

49. Wang Y, Kim JY, Park MS, Ji G. 2012. Novel *Bifidobacterium* promoters selected through microarray analysis lead to constitutive high-level gene expression. J. Microbiol. 50, 638–643.

50. Wang Z, Ye S, Li J, Zheng B, Bao M, Ning G. 2011. Fusion primer and nested integrated PCR (FPNI-PCR): a new high-efficiency strategy for rapid chromosome walking or flanking sequence cloning. BMC Biotechnol. 11, 109.

51. Wei C, Li M, Cao X, Jin Z, Zhang C, Xu M, Chen K, Zhang B. 2022. Linalool synthesis related *PpTPS1* and *PpTPS3* are activated by transcription factor *PpERF61* whose expression is associated with DNA methylation during peach fruit ripening. Plant Sci. 317, 111200.

52. Wei X, Geng M, Yuan J, et al. 2024. *GhRCD1* promotes cotton tolerance to cadmium by regulating the *GhbHLH12*-*GhMYB44*-*GhHMA1* transcriptional cascade. Plant Biotechnol. J. 22, 1777–1796.

53. Wu Z, Li L, Liu H, et al. 2021. *AaMYB15*, an R2R3-MYB TF in *Artemisia annua*, acts as a negative regulator of artemisinin biosynthesis. Plant Sci. 308, 110920.

54. Xu P, Su H, Jin R, Mao Y, Xu A, Cheng H, Wang Y, Meng Q. 2020. Shading effects on leaf color conversion and biosynthesis of the major secondary metabolites in the albino tea cultivar “Yujinxiang”. J. Agric. Food Chem. 68, 2528–2538.

55. Yan Y, Tao H, He J, Huang S. 2020. The HDOCK server for integrated protein-protein docking. Nat. Protoc. 15, 1829–1852.

56. Yang B, Lee M, Lin M, Chang W. 2022. 5-Azacytidine increases tanshinone production in *Salvia miltiorrhiza* hairy roots through epigenetic modulation. Sci. Rep. 12, 9349.

57. Yang J, Gu D, Wu S, Zhou X, Chen J, Liao Y, Zeng L, Yang Z. 2021. Feasible strategies for studying the involvement of DNA methylation and histone acetylation in the stress-induced formation of quality-related metabolites in tea (*Camellia sinensis*). Hortic. Res. 8, 253.

58. Yang L, Wen K, Ruan X, Zhao Y, Wei F, Wang Q. 2018. Response of plant secondary metabolites to environmental factors. Molecules 23, 762.

59. Zang Y, Xie L, Su J, Luo Z, Jia X, Ma X. 2023. Advances in DNA methylation and demethylation in medicinal plants: a review. Mol. Biol. Rep. 50, 7783–7796.

60. Zeng F, Li X, Qie R, Li L, Ma M, Zhan Y. 2020. Triterpenoid content and expression of triterpenoid biosynthetic genes in birch (*Betula platyphylla* Suk) treated with 5-azacytidine. J. For. Res. 31, 1843–1850.

61. Zha L, Liu S, Liu J, Jiang C, Yu S, Yuan Y, Yang J, Wang Y, Huang L. 2017. DNA methylation influences chlorogenic acid biosynthesis in *Lonicera japonica* by mediating *LjbZIP8* to regulate phenylalanine Ammonia-Lyase 2 expression. Front. Plant Sci. 8, 1178.

62. Zhang H, Lang Z, Zhu J. 2018. Dynamics and function of DNA methylation in plants. Nat. Rev. Mol. Cell Biol. 19, 489–506.

63. Zhang Y, Ji A, Xu Z, Luo H, Song J. 2019. The AP2/ERF transcription factor *SmERF128* positively regulates diterpenoid biosynthesis in *Salvia miltiorrhiza*. Plant Mol. Biol. 100, 83–93.

64. Zhao Q, Du Y, Wang H, Rogers HJ, Yu C, Liu W, Zhao M, Xie F. 2019. 5-Azacytidine promotes shoot regeneration during Agrobacterium-mediated soybean transformation. Plant Physiol. Biochem. 141, 40–50.

65. Zhao Y, Liu G, Yang, F, et al. 2023. Multilayered regulation of secondary metabolism in medicinal plants. Mol. Hortic. 3, 11.

66. Zhou P, Li H, Lin Y, et al. 2023. Omics analyses of *Rehmannia glutinosa* dedifferentiated and cambial meristematic cells reveal mechanisms of catalpol and indole alkaloid biosynthesis. BMC Plant Biol. 23, 463.

